# An efficient method for high molecular weight bacterial DNA extraction suitable for shotgun metagenomics from skin swabs

**DOI:** 10.1101/2023.02.23.529690

**Authors:** Iliana R. Serghiou, Dave Baker, Rhiannon Evans, J. Dalby Matthew, Raymond Kiu, Eleftheria Trampari, Sarah Phillips, Rachel Watt, Thomas Atkinson, Barry Murphy, Lindsay J. Hall, Mark A. Webber

## Abstract

The human skin microbiome represents a variety of complex microbial ecosystems that play a key role in host health. Molecular methods to study these communities have been developed but have been largely limited to low-throughput quantification and short amplicon sequencing, providing limited functional information about the communities present. Shotgun metagenomic sequencing has emerged as a preferred method for microbiome studies as it provides more comprehensive information about the species/strains present in a niche and the genes they encode. However, the relatively low bacterial biomass of skin, in comparison to other areas such as the gut microbiome, makes obtaining sufficient DNA for shotgun metagenomic sequencing challenging. Here we describe an optimised high-throughput method for extraction of high molecular weight DNA suitable for shotgun metagenomic sequencing. We validated the performance of the extraction method, and analysis pipeline on skin swabs collected from both adults and babies. The pipeline effectively characterised the bacterial skin microbiota with a cost and throughput suitable for larger longitudinal sets of samples. Application of this method will allow greater insights into community compositions and functional capabilities of the skin microbiome.

**Impact Statement:** Determining the functional capabilities of microbial communities within different human microbiomes is important to understand their impacts on health. Extraction of sufficient DNA is challenging, especially from low biomass samples, such as skin swabs suitable for shotgun metagenomics, which is needed for taxonomic resolution and functional information. Here we describe an optimised DNA extraction method that produces enough DNA from skin swabs, suitable for shotgun metagenomics, and demonstrate it can be used to effectively characterise the skin microbiota. This method will allow future studies to identify taxonomic and functional changes in the skin microbiota which is needed to develop interventions to improve and maintain skin health.

**Data Summary:** All sequence data and codes can be accessed at:

NCBI Bio Project ID: PRJNA937622

DOI: https://github.com/quadram-institute-bioscience/coronahit_guppy

DOI: https://github.com/ilianaserghiou/Serghiou-et-al.-2023-Codes

## Introduction

The skin microbiome is a complex ecosystem organised into distinct microbial communities present at different body sites (NASEM, 2018; Costello, et al., 2009). These microbial ecosystems participate in the host’s skin physiological functions and immunity (Cho and Blaser, 2012; Human Microbiome Project Consortium, 2012). Perturbations in these communities can negatively impact skin health, particularly early in life (Kong, 2011). Studying the skin microbiota and how it forms and changes over time is therefore important to understand how interventions that alter the microbiota affect skin health.

Previous skin microbiome studies have commonly used traditional 16S rRNA gene amplicon sequencing (metataxonomics) to taxonomically classify these complex communities (Jo, et al., 2016). This method is typically performed using the Illumina sequencing technology, which results in short reads for taxonomic classification to genus level (Pearman, et al., 2020). 16S rRNA gene amplicon sequencing provides limited taxonomic information on bacteria and archaea however does not tell us anything about strain variations or functional capacities; Alternatively, the use of Shotgun Metagenomic Sequencing (SMS) for taxonomic classification follows sequencing of all genetic material and is not limited to targeted regions (Sfriso, et al., 2020; Kuczynski, et al., 2012; Allaband, et al., 2019). This reduces bias from selective amplification efficiency and can provide taxonomic information at species/strain level as well as being able to provide information about functional capacities present in the microbiome and individual species (Jo, et al., 2016; Liu, et al., 2020; Sfriso, et al., 2020). SMS can be performed using multiple technologies, including the Illumina, Oxford Nanopore (ONT) and PacBio Single Molecule Real-Time (SMRT) platforms (Pearman, et al., 2020; Amarasinghe, et al., 2020). In contrast to the Illumina technology, the ONT and PacBio SMRT technologies produce long sequence reads. Data produced with these platforms will usually reconstruct more complete genomes than from short reads and facilitates the generation of high-quality Metagenome Assembled Genomes (MAGs) (Pearman, et al., 2020), which can be used for higher taxonomic resolution and functional information (Singleton, et al., 2021; Liu, et al., 2020).

The relatively low bacterial biomass of skin complicates the extraction of sufficient DNA quantities for SMS (Bjerre, et al., 2019; de Goffau, et al., 2018). This is particularly true for longer read technologies where more input material is needed (Wang, et al., 2021). There are a limited number of commercialised kit protocols available that can produce high molecular weight (HMW) DNA from skin in sufficient quantities for SMS, although none have been specifically optimised to extract DNA from skin microbiome samples. To address this need we describe here an optimised high-throughput automated DNA extraction method, for recovery of HMW microbial DNA from skin swabs. This was validated using skin swabs from adult volunteers and babies enrolled in the Pregnancy and Early Life (PEARL) study (Phillips, et al., 2021). The method results in DNA with yield and molecular weight suitable for SMS.

## Methods

### DNA extraction method development

To optimise extraction of microbial DNA from skin swabs, a Promega Maxwell® RSC 48 Instrument and RSC Blood DNA Kit (see Supplementary file 1 for protocol) were used as a starting point and different diluents and lysis procedures were evaluated for effectiveness. This instrument and kit were chosen as they produce HMW DNA (Mandrekar, et al., 2007, Bey, et al., 2010), with a higher binding capacity and cleaner eluate than traditional silica-based DNA purification systems (Sui, et al., 2020; Moeller, et al., 2014; Dunbar, et al., 2018; Promega, 2020). The platform also permits a high-throughput automated genomic DNA isolation from 48 samples in 40 minutes (Promega, 2020) making this system compatible with larger sample sets.

To obtain enough DNA from skin swabs, suitable for SMS, we optimised the RSC protocol by testing different variables including the initial diluent and various lysis procedures. After dilution and lysis, samples were heated, following the RSC Blood DNA Kit protocol, and loaded to the Maxwell instrument for the automated extraction (Figure 1).

**Figure 1.**
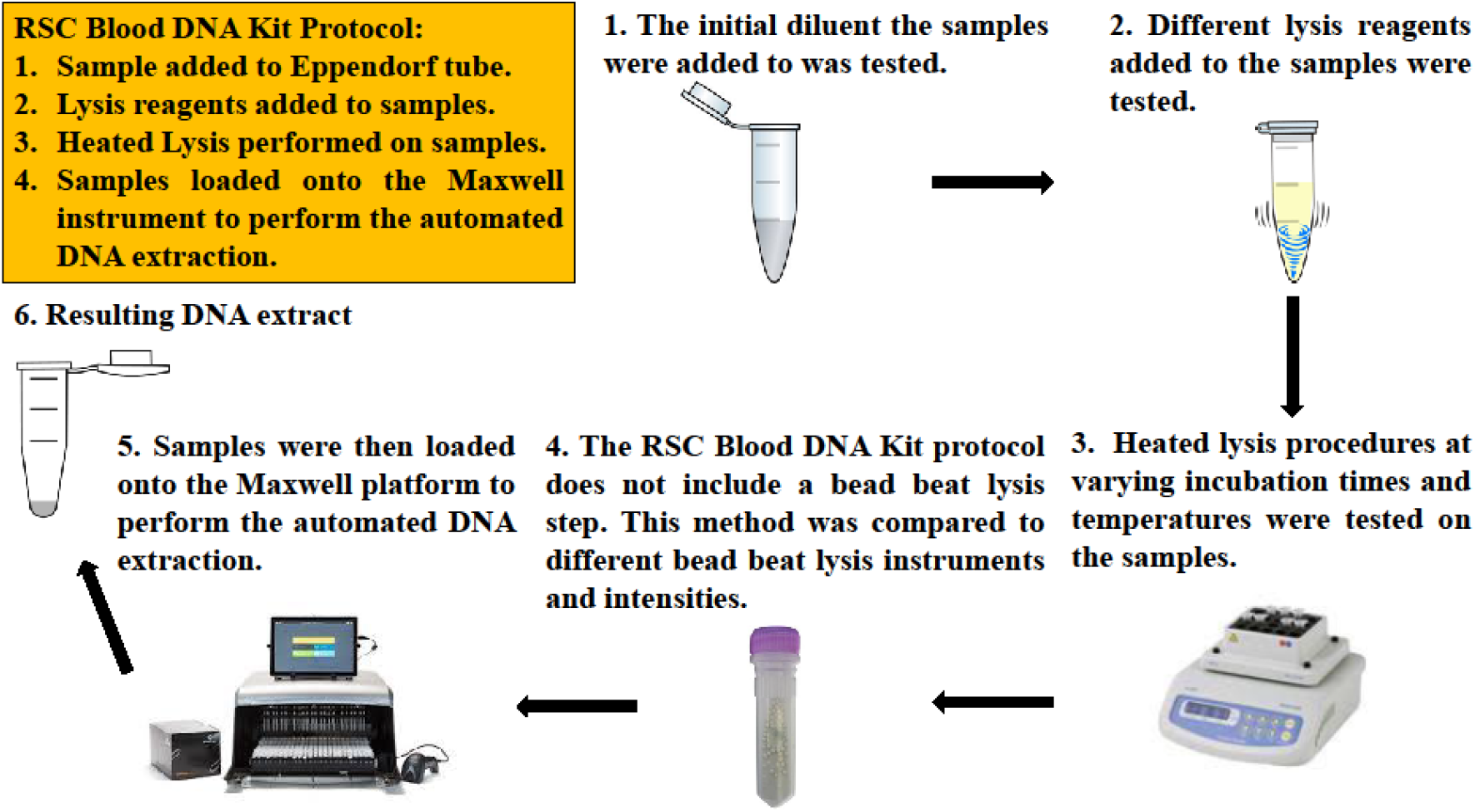
The RSC Blood DNA Kit protocol (yellow box) and alterations to test different initial diluents and lysis procedures.

#### Testing initial diluents: Measuring extracted bacterial DNA quantity and cellular viability

To allow for a protocol where a swab could be processed allowing both DNA extraction and, in parallel, culture of organisms, it was desirable to remove material from the swab into a diluent. To determine if diluents impacted bacterial viability and ability to extract DNA, 1x Phosphate Buffered Saline (PBS) and Milli-Q water, for collecting skin bacteria, were compared by measuring extracted bacterial DNA quantity recovered from swabs inoculated with bacteria. 44 sterile charcoal cotton swabs (M40-A2, Technical Service Consultants Ltd.) were used to collect a single colony from an agar plate inoculated with *Staphylococcus aureus* NCTC 8532 to act as a target for DNA extraction. These ‘spiked’ swab heads were snapped into 1.5ml Eppendorf tubes containing 1ml of either 1x PBS or Milli-Q water. These were then extracted following the Promega Maxwell® RSC 48 Instrument and RSC Blood DNA Kit protocol in Supplementary file 1,wih the following modification. The swabs were vortexed at full speed for 2 minutes and then centrifuged at 14,000 x g for 15 minutes to pellet the cells before the supernatant was removed, and cells were resuspended in 300µl of 1x PBS or Milli-Q water. Steps 4 and 6-8 of the RSC protocol were then followed. A bead beating step was then performed using a ‘FastPrep’ instrument for 3 minutes at setting 6.0. The samples were centrifuged again at 14,000 x g for 15 minutes to pellet the cells before sample supernatants were loaded onto the Maxwell instrument and the extraction started following steps 9-21 of the RSC protocol.

The effectiveness of 1x PBS and Milli-Q water, as initial diluents for collecting skin bacteria, was further compared by measuring bacterial cell viability through the recovery of bacteria from liquid cultures. Cell viability is an important factor as we wanted an initial dilution step which maintained bacterial viability and was therefore compatible with both culture of bacteria from samples and efficient DNA extraction. Overnight liquid cultures (10 ml) were grown from isolates of three species (*S. aureus* NCTC 8532, *Pseudomonas aeruginosa* PA14 and *Escherichia coli* EC18PR-0166-1, a food isolate of ST10), with three replicates for each. For each replicate, 1ml was transferred into a 15 ml falcon tube and pelleted by centrifugation at 14,000 x g for 15 minutes. Samples were then resuspended in 200µl of LB, 1x PBS or Milli-Q water and left for 1 hour at ambient temperature.

Serial dilutions of the resuspended samples were made and plated onto drug-free agar and incubated, which were then used to count viable numbers of cells in each sample. A total of nine independent samples were tested for each species in each diluent.

#### Testing lysis methods: Six extraction method procedures

Six lysis methods were compared to identify the best method for high yields of high molecular weight DNA from both Gram-negative and Gram-positive bacteria. Each method varied factors from common lysis methods used in commercial kits for research – heat, chemical, enzymatic, and mechanical (Gill, et al., 2016; Martzy, et al., 2019). Table 1 lists the differences between the six methods. Methods were tested using both overnight liquid cultures and sterile swab heads inoculated with harvested bacteria from overnight plate cultures.

**Table 1.** Comparison of extraction methods

|  | *Method 1* | *Method 2* | *Method 3* | *Method 4* | *Method 5* | *Method 6* |
| --- | --- | --- | --- | --- | --- | --- |
| **Heated lysis Step** | Yes | Yes | Yes | Yes | Yes | Yes |
| **Time** | 20 mins | 20 mins | 18h | 18h | 18h | 18h |
| **Temperature** | 56°C | 56°C | 37°C | 37°C | 37°C | 37°C |
| **Reagents** | Proteinase K, lysis buffer | Proteinase K, lysis buffer | Epicentre ready-lyse lysozyme | Epicentre ready-lyse lysozyme | Thermo Fischer lysozyme | Thermo Fischer lysozyme |
| **Agitation** | No | No | 300rpm | 300rpm | 300rpm | 300rpm |
| **Bead beat step** | Yes | Yes | Yes | Yes | Yes | Yes |
| **Instrument** | FastPrep | Tissue Lyser | FastPrep | Tissue Lyser | FastPrep | Tissue Lyser |
| **Settings** | 3 mins at 6.0 FastPrep | 3 mins at 20 Hz | 3 mins at 6.0 FastPrep | 3 mins at 20 Hz | 3 mins at 6.0 FastPrep | 3 mins at 20 Hz |
| **Heated Offboard Lysis Step** | No | No | Yes | Yes | Yes | Yes |
| **Temperature** | N/A | N/A | 68 °C | 68 °C | 68 °C | 68 °C |
| **Time** | N/A | N/A | 15 mins | 15 mins | 15 mins | 15 mins |
| **Reagents** | N/A | N/A | Proteinase K, buffer ATL, carrier RNA, buffer ACL | Proteinase K, buffer ATL, carrier RNA, buffer ACL | Proteinase K, buffer ATL, carrier RNA, buffer ACL | Proteinase K, buffer ATL, carrier RNA, buffer ACL |
| **Agitation** | N/A | N/A | 300rpm | 300rpm | 300rpm | 300rpm |

Duplicate 10 ml overnight liquid cultures were grown for each species (*S. aureus, P. aeruginosa* and *E. coli*), from each, 300µl was added into two 1.5ml Eppendorf tubes resulting in 6 tubes which were tested for method 1 and 2. A further 400µl of each liquid culture was added into four tubes resulting in 6 tubes tested for each remaining method. All samples were then extracted following the Promega Maxwell® RSC 48 Instrument and RSC Blood DNA Kit protocol (detailed in supplementary file 1) with changes to the lysis procedure for each of the six methods tested. All Eppendorf tubes were then vortexed at full speed for 2 minutes and centrifuged at 14,000 x g for 15 minutes to pellet the cells; the supernatants were removed, and pellets resuspended in 300µl (methods 1 or 2) or 400µl (methods 3-6) of 1x PBS.

For method 1 and 2 samples, 30µl of Proteinase K and 300µl of Lysis Buffer were added to the 300µl sample suspensions. These were then incubated in a heating block at 56°C for 20 minutes. For methods 3 and 4 samples, 3µl of Ready-Lyse lysozyme (Epicentre, 250U/µl in TES buffer) was added to the 400µl sample suspensions. For methods 5 and 6 samples, 3µl of Thermo Fischer lysozyme (250U/µl in TES buffer) was added to the 400µl sample suspensions. Samples from methods 3-6 were then incubated with agitation at 300rpm, 37°C for 18 hours. A bead beating step was performed on all samples. Method 1, 3 and 5 samples used the FastPrep instrument for 3 minutes at setting 6.0 and method 2, 4 and 6 samples used a Tissue Lyser instrument for 3 minutes at 20Hz to compare the impact of a less intense bead beating step. An off-board lysis was performed on method 3-6 samples, which included addition of 40µl proteinase K, 165µl Buffer ATL, 120µl Carrier RNA (lyophilised Carrier RNA was resuscitated with Buffer AVE to make a 1μg/µl solution), and 315µl Buffer ACL into the 400µl sample suspensions. These samples were then incubated at 68 °C for 15 minutes.

Samples from all methods were centrifuged at 14,000 x g for 15 minutes to pellet cells and the supernatants were loaded onto the Maxwell instrument and the extraction started following steps 9-21 of the initial RSC protocol.

After evaluation of the performance of the different methods from cultured cells, method 6 performed the best (see results) and was chosen for validation using swab samples. For validation, sterile charcoal cotton swabs (M40-A2, Technical Service Consultants Ltd.) were spiked with one colony from overnight plate cultures of each of the three species and eight independent swabs were processed per species. Swab heads were snapped off into 1.5ml Eppendorf tubes containing 1ml of 1x PBS and samples were vortexed for 2 minutes before being centrifuged at 14,000 x g for 15 minutes to pellet the cells. The supernatants were removed, and the pellets were resuspended with 400µl 1x PBS. The method 6 procedure was then followed as described above.

### Validation of DNA extraction method using volunteer and PEARL study skin swabs

The optimised DNA extraction method was tested on skin swabs from adults and babies to validate the selected method ability to obtain appropriate bacterial DNA for SMS and confirm data was suitable for analysing the taxonomic profiles of bacterial communities present on skin. Samples were cultured in parallel to DNA sequencing; this allowed us to identify organisms which should be represented in the SMS data whilst also enabling the creation of a skin microbiota culture collection for future functional work with strains of interest. Swabs were cultured aerobically and anaerobically on Columbia blood agar plates as in previous studies (Ogai, et al., 2018). For each swab, cells grown on the aerobic and anaerobic plates were harvested into one glycerol stock, a sample of which was then used for DNA extraction and SMS to compare to results direct from swabs.

#### Study design for adult volunteer and PEARL study baby skin swab collection

The Norwich Research Park Biorepository recruited and consented 12 adult volunteers between the age of 23-65. There was no contact between the researcher and participants to ensure anonymity. Eligible volunteer participants had no current skin conditions or had been prescribed antibiotics over the last 3 months. The volunteer participants were provided with Participant Information Sheets (PIS) and were consented with Consent Forms (CF) and provided samples using a self-swabbing protocol under observation and following instruction from Biorepository staff (Supplementary file 2). The volunteers collected two swabs, one from the right arm and one from the left arm, to produce 24 samples in total. Samples were stored in a 4°C fridge and anonymised with a unique barcode before being collected and tested on the same day swabbing was performed. In addition to the adult volunteers, swabs from the skin of ten babies collected at four months as part of the PEARL study were also included (see Phillips, et al., (2021) for study design and inclusion criteria, and Table S1 for baby participant metadata).

#### Volunteer and baby skin swab processing and finalised DNA extraction procedure

The skin swabs were processed as described above with the optimised method, a cell-free, diluent-only sample was included as a negative control on each extraction run and an established commercial mock community (the ATCC skin microbiome whole cell mix) was included as a positive control (ATCC, 2022). Dilutions of the positive control microbiome mix were also prepared to validate extraction efficiency and identify a cut-off point of starting material needed for SMS. For full details on the sample processing, DNA extraction protocol and the ATCC positive control protocol, see supplementary file 3.

### DNA quantification and quality assessment

A High Sensitivity (HS) assay using the Qubit 2.0 fluorometer instrument and HS Qubit Invitrogen kit, was used to quantify all samples. If a concentration was out of range, i.e., too high, the Broad Range (BR) Qubit assay was used instead, using the Qubit 2.0 fluorometer instrument and BR Qubit Invitrogen kit. Tapestation assays were used to determine DNA molecular weight. A D5000 or HS D5000 Tapesation assay were used with an Agilent 2200 instrument and Agilent D5000 or HS D5000 kits.

#### Shotgun Metagenomic Sequencing using Illumina and Oxford Nanopore

Preparation of libraries for SMS for both Illumina (Illumina DNA Prep Kit: 20018704) and ONT (Illumina® DNA Prep: 20018704, Tagmentation: 20060059) platforms included DNA normalisation, tagmentation, PCR barcoding, quantification, pooling, and quality control. Samples were then loaded onto the Illumina NextSeq500 Instrument using a Mid-output 300 cycle kit (Illumina Catalogue FC-404-2003) or the MinION flow cell ONT instrument (R9.4.1). The QIB Bioinformatics team converted the Illumina raw data to 8 FASTQ files for each sample, and the ONT raw data was converted into FASTQ files using the customised guppy method. All FASTQ files were then run through FastP (V.0.19.5+galaxy1) (Chen, et al., 2018), which is a pre-processing tool for FASTQ files that removes adaptors. For full details on the SMS protocol for Illumina and ONT, view Supplementary File 4.

#### Generating taxonomic profiles

All SMS data was automatically deposited in a local instance of IRIDA (irida-19.09.2) (Matthews, et al., 2018) and uploaded to the QIB Galaxy platform (V.19.05) (Afgan, et al., 2018). Here, data was cleaned by removing adaptors and trimming reads, and filtered for quality using Fastp (V.0.20.0) (-q 20) (Chen, et al., 2018), before reads mapping against a human reference database (human_20200311) were removed using Kraken2 (V.2.1.1+galaxy0) (Wood, et al., 2019). Remaining reads were then analysed to obtain microbiota taxonomic profiles using Kraken2 (V.2.1.1+galaxy0) (Wood, et al., 2019) and Bracken (V.2.2) (Lu, et al., 2017).

#### MAG extraction

Using the trimmed and filtered reads, host-associated sequences were removed via Kneaddata (V.0.10.0) (The Huttenhower Lab) with human genome (GRCh38.p13) to generate clean fastq reads. Shotgun metagenome raw reads were co-assembled with MEGAHIT (V.1.2.9) (Li, et al., 2015) prior to extraction of MAGs. The MetaWRAP (V.1.3.2) pipeline (Uritskiy, et al., 2018) was used to extract MAGs based upon metagenome assemblies generated and metagenome clean reads via binning software ‘metaBAT’ (V.2.12.1) (Kang, et al., 2015), ‘MAXBIN2’ (V.2.2.6) (Wu, et al., 2016) and ‘CONCOCT’ (V.1.1.0) (Alneberg, et al., 2013) using the sub-module ‘binning’. MAGs were then refined using sub-module ‘bin_refinement’ to select the high-quality bins from each sample with completeness >80% and contamination <10% according to CheckM (V.1.1.3) (Parks, et al., 2015). All MAGs were taxonomically ranked using gtdb-tk (V.1.5.1) (Chaumeil, et al., 2020) via module gtdbtk classify_wf.

#### Data visualisation

R (V.4.1.2) (RStudio Team, 2021) and the package ggplot2 (Wickham, 2009) were used to plot taxonomic profiles and alluvial and box plots. GraphPad Prism (V.5.04) (GraphPad Software, 2010) was used to generate scatter plots.

### Statistical Analysis

Statistical analysis was performed using Unpaired T-tests in GraphPad Prism (V.5.04) (GraphPad Software, 2010). A significance level of 0.05 was used to identify results likely to be different.

## Results

### Optimisation of DNA extraction method

#### Impact of initial diluents on extracted bacterial DNA quantity and cell viability

There was no significant difference between amounts of bacterial DNA extracted from the 44 sterile charcoal cotton swabs spiked with *S. aureus* and processed in either PBS or water (Figure 2A). Recovery of *S. aureus, P aeruginosa* and *E. coli*, also showed no significant differences in viable numbers recovered after suspension in either diluent (P > 0.05; Figure S1). As there was no significant difference in both DNA extraction and bacterial recovery between PBS and water, future experiments used PBS.

**Figure 2.**
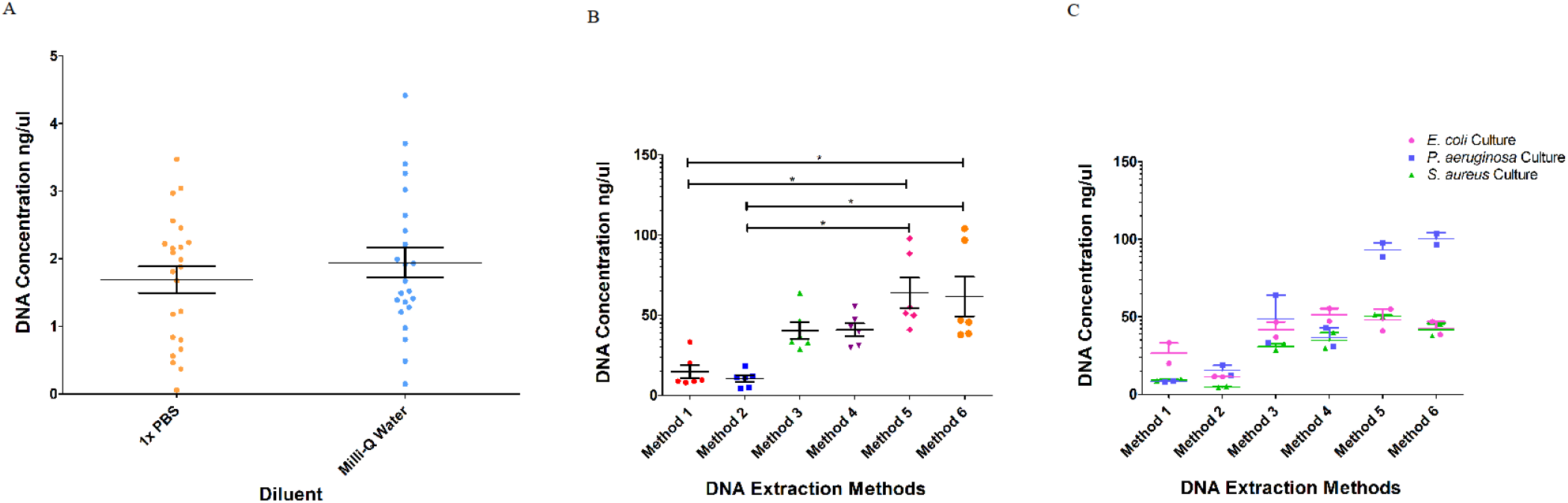
Results of the variables tested. A: Total DNA yield (ng/µl) from spiked swabs processed in 1xPBS and Milli-Q water during a DNA extraction. B: Total DNA yield (ng/µl) obtained from each DNA extraction method. C: Total DNA yield (ng/µl) per species for each method. Horizontal bars on each plot show averages, vertical bars show the standard error of the mean (SEM) and lines with an asterisk (*) indicate significant (p <0.05) differences.

#### Testing lysis methods: Six extraction method procedures

DNA extracted from liquid cultures of *S. aureus, P. aeruginosa* and *E. coli* using the six methods (Table 1), showed that methods 5 and 6 yielded the most DNA, (40.9-97.7ng/µl and 37-104ng/µl respectively), and there was a significant difference in DNA concentrations between methods 5 and 6 and other methods (Table S2; Figure 2B and 2C). There was no significant difference in extraction efficiency between each bacterial species. DNA extraction methods 2, 4 and 6 produced higher molecular weights than the others, ranging from 20232-31786 bp (Table 2). Together, these results demonstrated that method 6 produced the most DNA of highest molecular weight. This method was also the most cost effective due to the cheaper lysozyme used and was chosen for further validation. This method included overnight lysis with lysozyme, a further heated offboard lysis step and a bead beating lysis using a Tissue Lyser.

**Table 2.** Average molecular weight (bp) of DNA extracted

| Sample | Method 1 | Method 2 | Method 3 | Method 4 | Method 5 | Method 6 |
| --- | --- | --- | --- | --- | --- | --- |
| *E. coli* | 0 | 25321 | 1735 | 21693 | 1909 | 24219 |
| *E. coli* | 0 | 25697 | 2057 | 20232 | 1877 | 22730 |
| *P. aeruginosa* | 0 | 23771 | 1763 | 23163 | 1678 | 29459 |
| *P. aeruginosa* | 0 | 23826 | 1631 | 23742 | 1538 | 31786 |
| *S. aureus* | 0 | 25085 | 1831 | 23954 | 1668 | 22118 |
| *S. aureus* | 0 | 22011 | 1762 | 24084 | 1711 | 22904 |

DNA extractions from sterile charcoal cotton swabs spiked with independent cultures were successful, with DNA concentrations averaged at 22.1ng/µl.

### DNA extraction method validation using swabs from volunteers or babies

DNA concentrations from adult and baby skin swabs, that were extracted using method 6, ranged from < 0.50 (no detected DNA) – 10.5 ng/µl (Table S3) with DNA successfully extracted from all the baby samples but only 15/24 adult volunteer samples. Cultured plates recovered bacteria from all adult skin swabs although recovery of cultures from the baby samples was only successful for 4/10 swabs. Concentrations of DNA extracted from cultured bacteria averaged at 79.9ng/µl.

As some swabs did not yield DNA using method 6, we compared DNA yield from the extracted swabs after different overnight lysis incubation times. Samples were randomly incubated for either 18, 20 or 22 hours (Figure 3). A significant difference between 18 and 20 hours and 18 and 22 hours (P < 0.05) was observed, but no significant difference between 20 and 22 hours (P > 0.05). Samples that did not yield detectable amounts of DNA were those incubated for 18 hours therefore, future samples were incubated between 20-22 hours to obtain higher DNA yield.

**Figure 3.**
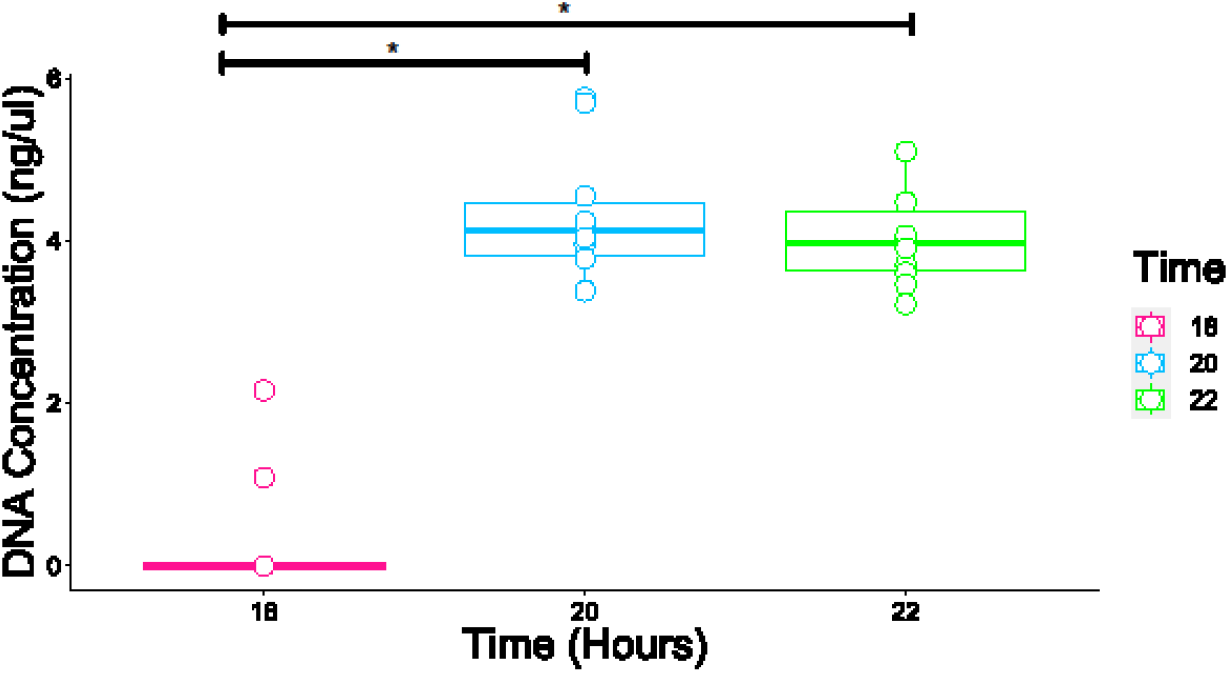
Comparison of DNA yield (ng/µl) from samples incubated for different periods. The box plots show the average DNA concentrations (ng/µl) for each incubation time. Horizontal bars on each plot show averages, vertical bars show the standard error of the mean (SEM) and lines with an asterisk (*) indicate significant (p <0.05) differences.

After removal of human reads, microbial taxonomic profiles were generated using both Illumina and ONT sequence data using Kraken2 and Bracken (Figures 4-6; swabs and cultures).

**Figure 4.**
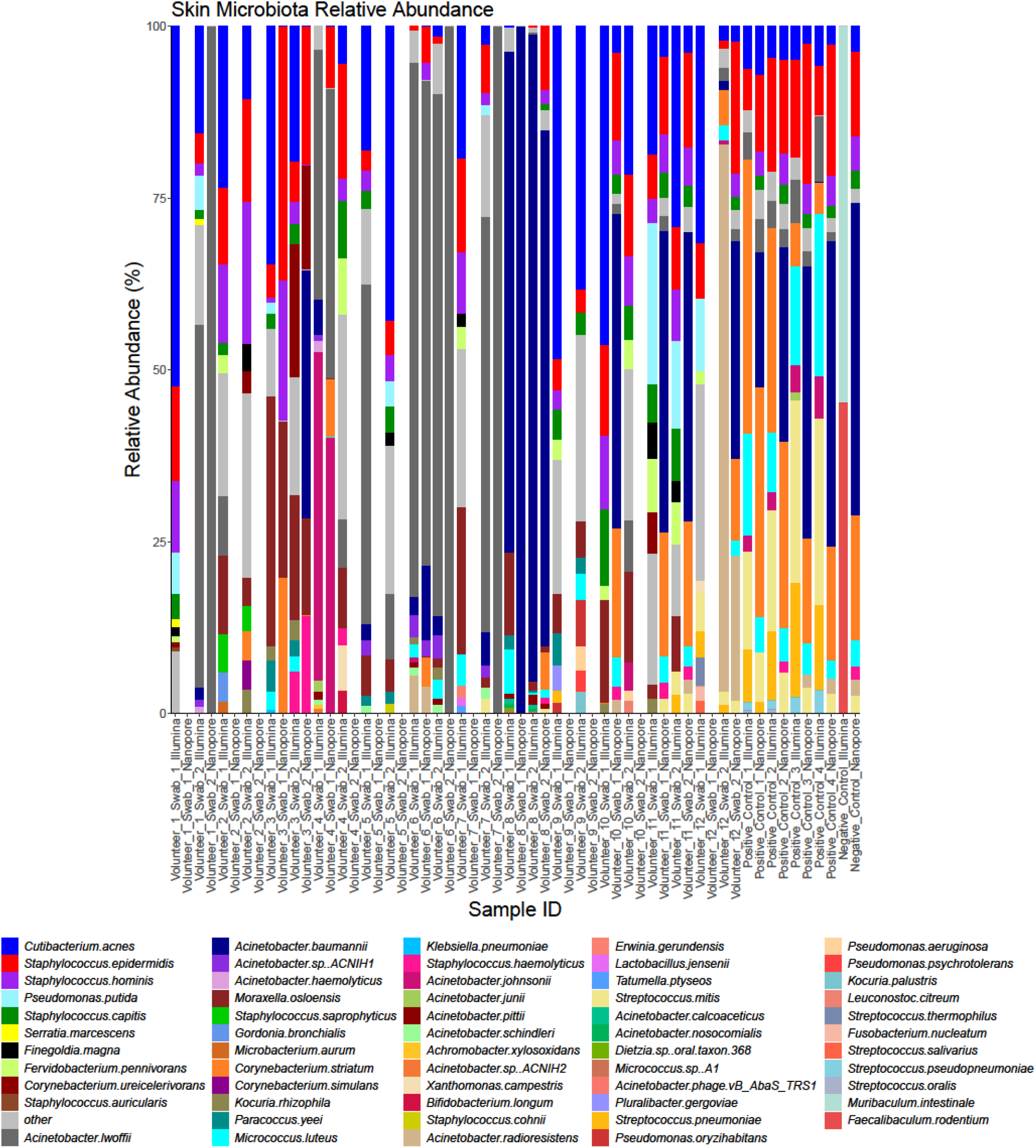
Taxonomic profiles of skin swab microbiota from 12 adult volunteers (two swabs collected from both forearms from each volunteer) generated using Illumina and Nanopore data. Profiles show the relative abundance (%) of the 10 most abundant species that occur within each sample.

**Figure 5.**
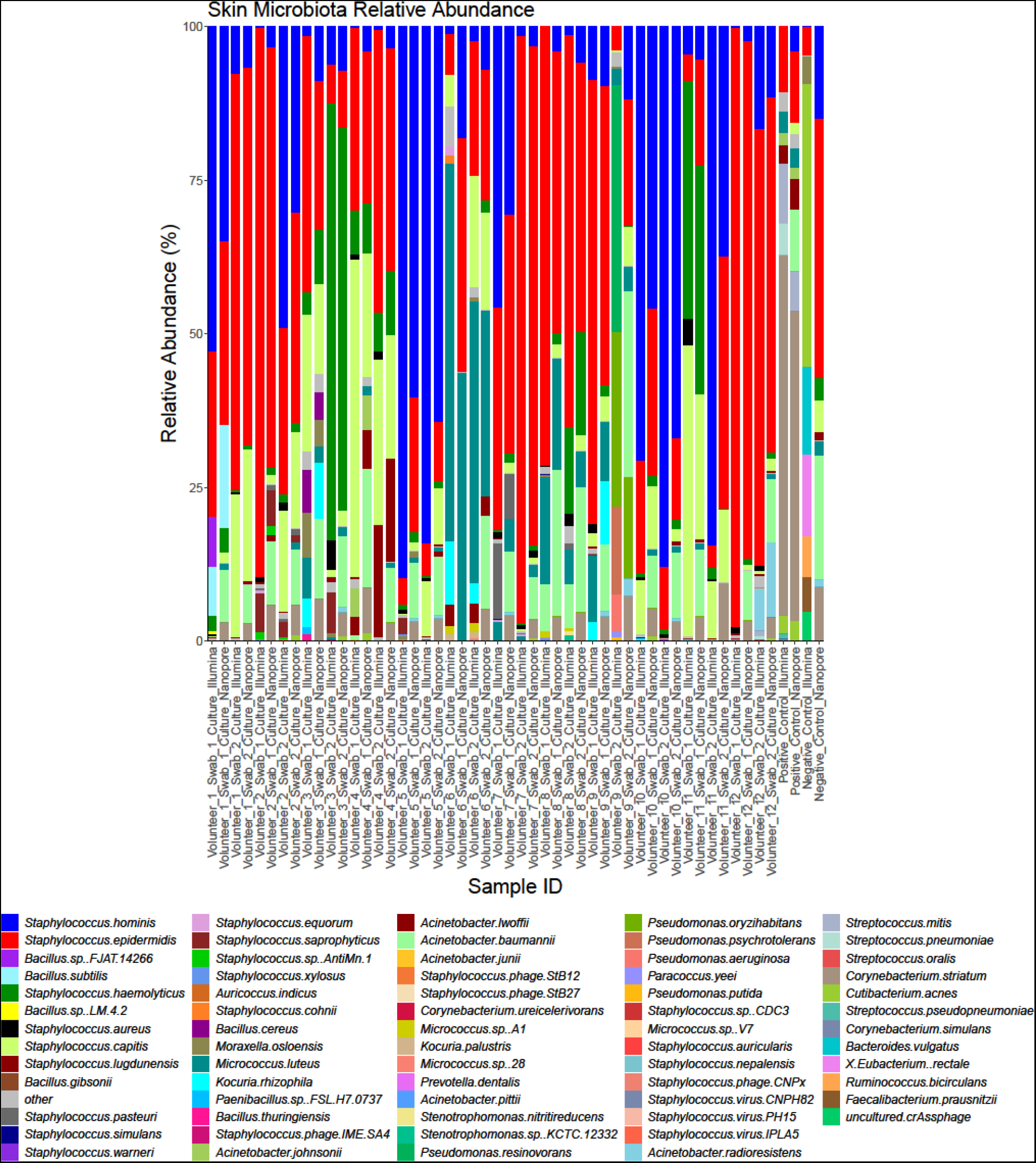
Taxonomic profiles of skin swab culture microbiota from 12 adult volunteers (two swabs collected from both forearms from each volunteer) generated using Illumina and Nanopore data. Profiles show the relative abundance (%) of the 10 most abundant species that occur within each sample.

**Figure 6.**
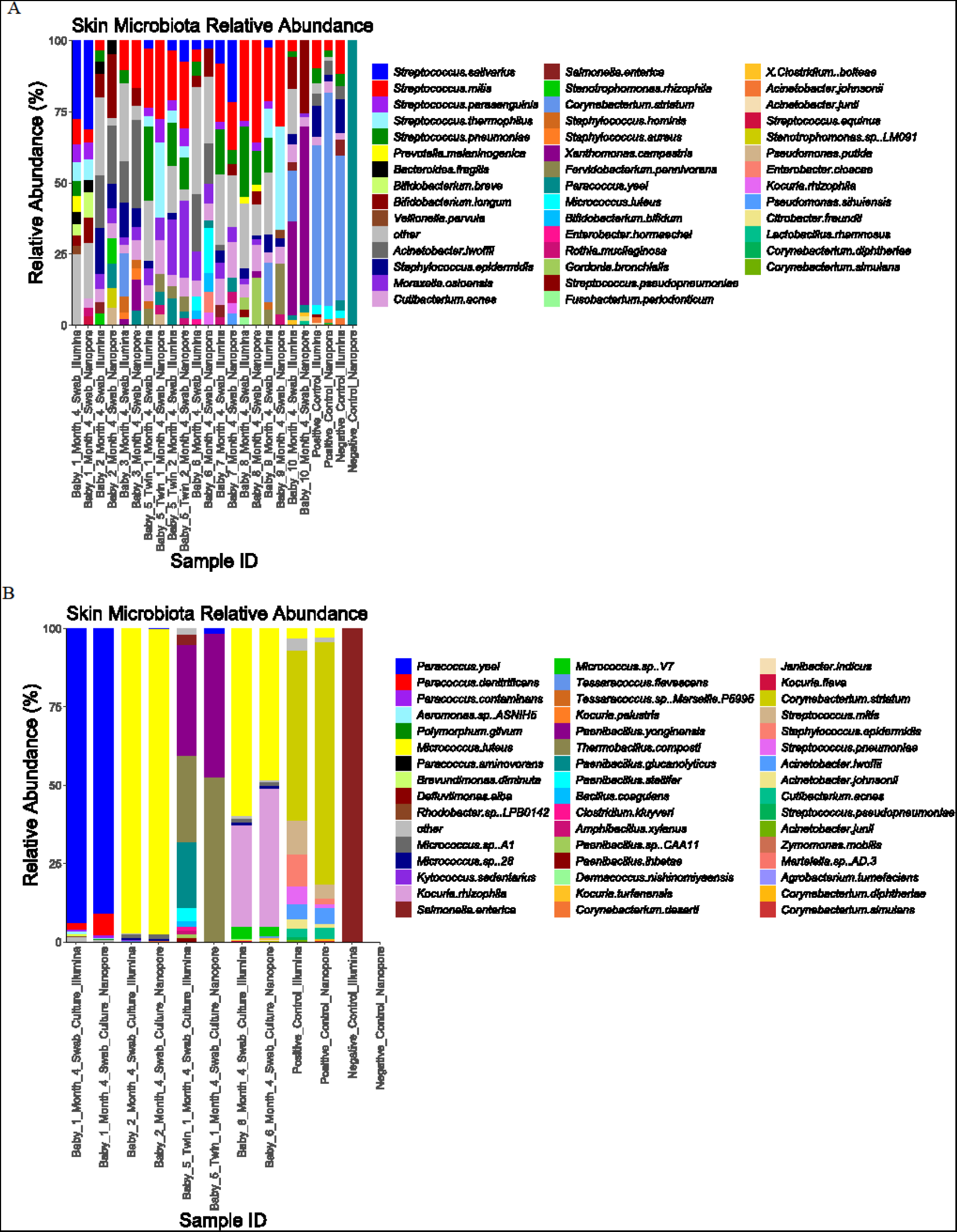
Taxonomic profiles of the skin swab and culture microbiota from ten PEARL babies (one swab collected off one forearm from each baby) generated using Illumina and Nanopore data. Profiles show the relative abundance (%) of the 10 most abundant species that occur within each sample. A: Illumina and Nanopore skin swab data. B: Illumina and Nanopore skin culture data.

The positive controls displayed the expected microbiota from the ATCC skin microbiome whole cell mix - *Acinetobacter johnsonii, Corynebacterium striatum, Micrococcus luteus, Cutibacterium acnes, Staphylococcus epidermidis, Streptococcus mitis*. There was also a clear reduction in reads from these samples across the dilution series. This demonstrated that the DNA extraction method was able to effectively extract DNA from the diverse range of species present in the ATCC skin microbiome mix. Some background contamination was detected in negative controls, although the number of reads was always much lower than in samples and most DNA fragments identified in the negative controls mapped against organisms not seen in the test samples.

Both Illumina and ONT data indicated a typical skin microbiota from both adult and baby skin swabs and generated enough reads for downstream taxonomic analysis at the species level. The adult swabs identified bacteria, viruses, and phages, whereas the baby swabs only displayed bacterial diversity.

Baby skin swabs contained more *Streptococcus* and fewer *Staphylococcus* species when compared to adult skin swabs. The baby skin swabs also indicated the presence of *Bifidobacterium longum, Bifidobacterium breve* and *Bifidobacterium bifidum*, which are not typical skin residents, but common residents of the infant gut, which likely demonstrates transient skin contamination on the babies (Toscano, et al., 2017; Yan, et al., 2021). Importantly, data was generated for skin swabs that had very low DNA concentrations. Illumina and ONT platforms identified very similar microbiota profiles for both skin swabs and cultures, with comparable percentage total counts of the most abundant species (those representing more than 0.5% of each sample) (Figure S2). Analysis of the taxonomic profile from cultured samples exhibited less microbial diversity than the skin swabs as expected but confirmed the presence of species identified in the SMS. As in the SMS data, adult cultures exhibited more *Staphylococcus* species than *Streptococcus*.

Once a successful DNA extraction method was established, the depth of sequence data required to provide optimal phylogenetic resolution and to construct MAGs were both assessed. This was done by comparing outcomes using 5Gbp per sample and subsamples thereof down to 1Gbp of data. For species identification a rarefaction curve was produced, which showed more species identified as more data was used; though statistical analysis showed there was not a significant difference in species recovery between 2.5 and 5Gbp of data (Figure S3A). Recovery of MAGS was also higher from samples where 5Gbp of data were used than 1Gbp, although this difference was not found to be statistically significant (Figure S3B; Table S4). Based on this analysis, 5Gbp of data appears to be adequate for phylogenetic analysis of the skin microbiota using this method, whilst also providing useful functional information.

## Discussion

We aimed to develop an efficient protocol for DNA extraction suitable for use from both skin swabs and cultured bacterial cells. Initial testing showed both water and PBS were suitable diluents to maintain viability and for DNA extraction in agreement with previous studies (Banning, et al., 2002; Liao and Shollenberger, 2003; Downey, et al., 2012) and PBS was then used throughout. Comparison of a variety of lysis procedures identified the effectiveness of a combined approach using both overnight heated enzymatic (lysozyme) and mechanical (bead beat) lysis methods to result in sufficient DNA yield of a high molecular weight from both Gram-positive and Gram-negative bacteria. Previous work has indicated that the type of enzyme and mechanical intensity is also important for lysis of different bacterial species (Schindler and Schuhardt, 1964; Yuan, et al., 2012; Albertesen, et al., 2015); however, our combined use of a mechanical and enzymatic lysis approach resulted in an unbiased extraction of Gram-positive and Gram-negative bacteria, which was validated by the production of expected profiles from the positive control mock community (Maghini, et al., 2021).

Given the low biomass of skin microbiota, some of the adult skin swabs produced very low/absent DNA concentrations and paired cultures also indicated low bacterial burden. Individual variations when swabbing (pressure, direction, frequency) can affect the yield of DNA and viable bacteria, and are difficult to control (Van Horn, et al., 2008) and may be responsible for this variation. A swabbing method was used as it is commonly used to collect skin microbiome samples (Van Horn, et al., 2008) and was already used by our local PEARL study to collect samples due to its non-invasive nature, which is suitable for neonates, who have an underdeveloped skin structure (Narendran, et al., 2010; Chiou and Blume-Peytavi, 2004). We also found a difference in sensitivity between platforms for samples with low amounts of DNA, some adult swabs did not produce data using the ONT platform although these same samples generated bacterial cultures. As the ONT platform requires more input DNA to generate data than Illumina platforms (Wang, et al., 2021), the inability to generate data for some samples was not surprising as skin swabs can be low biomass (Bjerre, et al., 2019; de Goffau, et al., 2018). However, increasing the overnight incubation time did improve DNA yield, and the Illumina sequencing resulted in generated data for all samples.

Most samples did generate data from both Illumina and ONT platforms which presented similar microbiota profiles from skin swabs and cultures. Typical adult skin microbiota (Phyla; *Pseudomonadota, Actinomycetota*, and *Bacillota*) (Grice, et al., 2009; Costello, et al., 2009; Byrd, et al., 2018) and infant skin microbiota (Phyla; *Bacillota, Actinomycetota, Pseudomonadota*, and *Bacteroidota)* (Capone, et al., 2011) were detected. We focused on bacterial species identified, but the protocol did identify other skin microbiota (viruses, phages and fungi), although only from adult volunteers (Byrd, et al., 2018). Other researchers can use this protocol as a starting point to be adapted if these organisms are their focus. Baby profiles only contained bacteria, and demonstrated less microbial diversity than adults, which has been shown in previous studies (Zhu, et al., 2019). Baby skin did exhibit more *Streptococcus* species than adult skin, which agrees with previous work demonstrating a predominance of *Streptococcus* species in early age, which decreases with age (Capone et al., 2011; Zhu, et al., 2019). Interestingly, sequencing of swabs from infant skin identified *Bifidobacterium* species, which are not typical skin residents, but rather maternal and infant gut residents and they can also be found in breast milk (Yan, et al., 2021) (Toscano, et al., 2017). Given the paired cultures did not result in any *Bifidobacterium* isolates, this is likely to indicate transient transfer to the babies’ skin through breast feeding. The babies with available metadata that showed *Bifidobacterium* presence on the skin were all breast fed at some point between birth and month 4.

Whilst skin is a relatively low biomass environment, we did not need to include any methods to mechanically deplete human DNA or selectively enrich microbial DNA before SMS (Marquet, et al., 2022), which have been needed in some other studies on low biomass samples. These enrichment approaches do not reliably target all species (Marquet, et al., 2022), can skew the resulting genomic profiles (Hammond, et al., 2016) and depletion can result in some loss of bacteria (Marquet, et al., 2022), thus further steps are required for downstream analysis. In our described method, we generated enough data, and depleted human DNA computationally, therefore precluding the need for any additional steps that may introduce biases and skew skin microbiota profiles.

Both Illumina and ONT sequence data allowed identification of all ATCC positive control species, with a clear reduction in read number across the dilution series. These results further demonstrate the effectiveness of the extraction method and utility of both sequencing platforms. Inclusion of a commercially available mixed community positive control, with a known cell concentration, is important for standardising the extraction process, and serial diluting the positive control can determine the limit of detection (Eisenhofer, et al., 2019). This is also helpful when comparing different sequencing runs and sample sets, allowing more robust comparisons to be made. Although, we tried to define a limit of detection for DNA concentration and read number required for effective SMS, we had several swab samples that did not obtain a DNA concentration reading, but usable reads were produced for taxonomic profiling. Therefore, no obvious cut-off for a limit of detection was determined, and indeed there is also no ‘defined’ limit identified in the literature for low biomass samples, such as skin swabs.

We did identify some background contamination in the negative controls, contamination commonly occurs in metagenomic studies, especially those with low biomass samples (Lou, et al., 2022). Several studies have identified contamination sources occurring from neighbouring samples and the ‘kitome’ (Lou, et al., 2022; Olomu, et al., 2020). Contamination within a dataset can be identified and removed using bioinformatic techniques (Zhou, et al., 2014; Davis, et al., 2018) although low biomass samples have a higher risk of true microbial microbiota members being removed (Diaz, et al., 2021). Given the background contaminants in the controls were at a very low level and mostly represented species not seen in the test samples we did not remove them as they had a negligible impact on the profiles produced.

We determined that the generation of 5Gbp of Illumina data from a skin swab was suitable for microbial species profiling but produced a limited number of MAGs. MAGs are important for in-depth functional information (Singleton, et al., 2021) and indicate genome quality (Bowers, et al., 2017; Parks, et al., 2015; Sczyrba, et al., 2017), and they can be used to identify novel taxa and allow further comparison with whole genome sequence data from isolates. Our method is compatible with both Illumina and ONT platforms and combining a higher sequencing depth with ONT data has potential to improve the number and quality of MAGs to be recovered (DeMaere and Darling, 2019; Gweon, et al., 2019; Singleton, et al., 2021).

## Abbreviations

PEARL: Pregnancy and Early Life Study
SMS: Shotgun Metagenomic Sequencing
ONT: Oxford Nanopore Technology
HMW: High Molecular Weight
LMW: Low Molecular Weight
PBS: Phosphate Buffered Saline
LB: Lysogeny Broth
MAGs: Metagenome Assembled Genomes
PIS: Participant Information Sheets
CF: Consent Forms
HS: High Sensitivity
BR: Broad Range
QIB: Quadram Institute of Biosciences

## Conclusion

An optimised medium-throughput DNA extraction, SMS, and analysis approach can effectively characterise the skin microbiota from adults and babies. This method can be applied for in-depth analysis of cohort studies allowing identification of taxonomic and functional changes of mothers and infants over time and should allow comparison to other body sites (e.g., the gut). Robust microbiota profiling, particularly in less well studied niches such as the skin, is important for the development of methods to alter microbiome compositions for health.

## Ethics

Ethical approval was obtained for the adult volunteer recruitment, skin swab sampling and processing from the University of East Anglia (UEA) Faculty of Medicine and Health Sciences (FHM); The recruitment, sampling and processing was performed under the Norfolk and Norwich University Hospital (NNUH) Biorepository ethics - FMH ethical approval reference: 2020/21-065.

The PEARL study was approved by The Quadram Institute Biosciences (QIB) Human Research Governance Committee (HRGC), local Research Ethics Committee (REC), and Health Research Authority (HRA). This study was conducted in accordance with the principles of the Declaration of Helsinki. The proposed research was conducted in accordance with the conditions and principles of the International Conference on Harmonisation Good Clinical Practice (ICH GCP), and in compliance with the UK national law. The research meets the requirements of the EU General Data Protection Regulation (GDPR), UK Data Protection Act 2018 and relevant sponsor’s policies - IRAS number: 241880.

## Author contributions

Conceptualisation, I.R.S., B.M., M.A.W., L.J.H.; Data curation, I.R.S., B.M., M.A.W., L.J.H., M.D., R.K., D.B., R.E., S.P., R.W. and T.A.; Formal analysis, I.R.S., B.M., M.A.W., L.J.H., M.D. and R.K.; Funding acquisition, B.M., M.A.W. and L.J.H.; Investigation, I.R.S., B.M., M.A.W., L.J.H., D.B., R.E., S.P., R.W. and T.A.; Methodology, I.R.S., B.M., M.W., L.J.H., D.B. and R.E.; Project administration, I.R.S., B.M., M.A.W., L.J.H., S.P., R.W. and T.A.; Resources, I.R.S., B.M., M.A.W., L.J.H., D.B., R.E., M.D., R.K., S.P., R.W., T.A.; Supervision, I.R.S., B.M., M.A.W. and L.J.H.; Validation, I.R.S., D.B., R.E., M.D. and R.K.; Visualisation, I.R.S., M.D. and R.K.; Writing – original draft, I.R.S., B.M., M.A.W., L.J.H., D.B., R.E., S.P., R.W., T.A., M.D., and R.K.; Writing – review and editing, I.R.S., E.T., B.M., M.A.W. and L.J.H.

## References

1. Afgan, E., Baker, D., Batut, B., van den Beek, M., Bouvier, D., Cech, M., Chilton, J., Clements, D., Coraor, N., Grüning, B. A., Guerler, A., Hillman-Jackson, J., Hiltemann, S., Jalili, V., Rasche, H., Soranzo, N., Goecks, J., Taylor, J., Nekrutenko, A., & Blankenberg, D. (2018). The Galaxy platform for accessible, reproducible and collaborative biomedical analyses: 2018 update. Nucleic acids research, Vol. 46(W1), W537–W544. https://doi.org/10.1093/nar/gky379

2. Albertsen M, Karst SM, Ziegler AS, Kirkegaard RH, Nielsen PH (2015). Back to Basics –The Influence of DNA Extraction and Primer Choice on Phylogenetic Analysis of Activated Sludge Communities. PLoS ONE, Vol. 10(7): e0132783. https://doi.org/10.1371/journal.pone.0132783

3. Allaband C, McDonald D., Vázquez-Baeza Y., Minich J.J., Tripathi A., Brenner D.A., Loomba R., Smarr L., Sandborn W.J. and Schnabl B., Dorrestein, P., Zarrinpar, A. and Knight, R., (2019). Microbiome 101: Studying, analyzing, and interpreting gut microbiome data for clinicians. Clinical Gastroenterology Hepatology, Vol. 17, pp. 218–230. https://doi.org/10.1016/j.cgh.2018.09.017

4. Alneberg, J., Bjarnason, B.S., de Bruijn, I., Schirmer, M., Quick, J., Ijaz, U.Z., Loman, N.J., Andersson, A.F., Quince, C., (2013). CONCOCT: Clustering cONtigs on COverage and ComposiTion. arXiv:1312.4038. https://doi.org/10.48550/arXiv.1312.4038

5. Amarasinghe S.L., Su S., Dong X., Zappia L., Ritchie M.E. and Gouil Q., (2020). Opportunities and challenges in long-read sequencing data analysis. Genome Biology, Vol. Vol. 21(30). https://doi.org/10.1186/s13059-020-1935-5

6. ATCC Skin Microbiome Whole Cell Mix. https://www.atcc.org/products/msa-2005 Accessed December 2022.

7. Banning, N., Toze, S. and Mee, B.J., (2002). Escherichia coli survival in groundwater and effluent measured using a combination of propidium iodide and the green fluorescent protein. Journal of Applied Microbiology, Vol. 93(1), pp. 69–76. https://doi.org/10.1046/j.1365-2672.2002.01670.x

8. Bey S.B., Fichot E.B., Dayama G., Decho A.W. and Norman R.S., (2010). Extraction of High Molecular Weight DNA from Microbial Mats. BioTechniques, Vol. 49(3): pp. 631–640. https://doi.org/10.2144/000113486

9. Bjerre R.D., Hugerth L.W., Boulund F., Seifert M., Johansen J.D. and Engstrand L., (2019). Effects of sampling strategy and DNA extraction on human skin microbiome investigations. Sci. Rep., Vol. 9, pp. 1–11. https://doi.org/10.1038/s41598-019-53599-z

10. Bowers, R., Kyrpides, N., Stepanauskas, R., et al., (2017). Minimum information about a single amplified genome (MISAG) and a metagenome-assembled genome (MIMAG) of bacteria and archaea. Nature Biotechnology, Vol. 35, pp. 725–731. https://doi.org/10.1038/nbt.3893

11. Byrd A.L., Belkaid Y. and Segre J.A., (2018). The human skin microbiome. Nature Reviews Microbiology, Vol. 16, 143–55. https://doi.org/10.1038/nrmicro.2017.157

12. Capone K.A., Dowd S.E., Stamatas G.N. and Nikolovski J., (2011). Diversity of the human skin microbiome early in life. Journal of Investigative Dermatology, Vol. 131(10), pp. 2026–https://doi.org/10.1038/jid.2011.168

13. Chaumeil, P-A., Mussig, A.J., Hugenholtz, P., Parks, D.H., (2020). GTDB-Tk: a toolkit to classify genomes with the Genome Taxonomy Database, Bioinformatics, Vol. 36(6), pp. 1925–1927. https://doi.org/10.1093/bioinformatics/btz848

14. Chen. S., Zhou, Y., Chen, Y. and Gu, J., (2018). Fastp: an ultra-fast all-in-one FASTQ preprocessor. Bioinformatics, Vol. 34(17), pp. i884–i890. https://doi.org/10.1093/bioinformatics/bty560

15. Chiou Y.B. and Blume-Peytavi U., (2004). Stratum corneum maturation. A review of neonatal skin function. Skin Pharmacology Physiology, Vol. 17(2), pp. 57–66. https://doi.org/10.1159/000076015

16. Cho I. and Blaser M.J., (2012). The human microbiome: At the interface of health and disease. Nature Reviews Genetics, Vol. 13, 260. https://doi.org/10.1038/nrg3182

17. Costello E.K., Lauber C.L., Hamady M., Fierer N., Gordon J.I. and Knight R., (2009). Bacterial community variation in human body habitats across space and time. Science, Vol. 326(5960), pp. 1694–1697. https://doi.org/10.1126/science.1177486

18. Davis, N.M., Proctor, D.M., Holmes, S.P., Relman, D.A. and Callahan, B.J., (2018). Simple statistical identification and removal of contaminant sequences in marker-gene and metagenomics data. Microbiome, Vol. 6(226). https://doi.org/10.1186/s40168-018-0605-2

19. de Goffau, M.C., Lager, S., Salter, S.J., Wagner, J., Kronbichler, A., Charnock-Jones, D.S., Peacock, S.J., Smith, G.C.S. and Parkhill, J., (2018). Recognizing the reagent microbiome. Nature Microbiology, Vol. 3, pp. 851–853. https://doi.org/10.1038/s41564-018-0202-y

20. DeMaere, M.Z. and Darling, A.E., (2019). bin3C: exploiting Hi-C sequencing data to accurately resolve metagenome-assembled genomes. Genome Biology, Vol. 20(46). https://doi.org/10.1186/s13059-019-1643-1

21. Díaz, S., Escobar, J.S. and Avilaa, F.W., (2021). Identification and Removal of Potential Contaminants in 16S rRNA Gene Sequence Data Sets from Low-Microbial-Biomass Samples: an Example from Mosquito Tissues. American Society for Microbiology, Vol. 6(3), pp. e00506–21. https://doi.org/10.1128/mSphere.00506-21

22. Downey, A.S., Da Silva, S.M., Olson N.D., Filliben, J.J., Morrowa, J.B., (2012). Impact of Processing Method on Recovery of Bacteria from Wipes Used in Biological Surface Sampling. Applied and Environmental Microbiology, Vol. 78(16), pp. 5872–5881. https://doi.org/10.1128/AEM.00873-12

23. Dunbar J., Gallegos-Gravesa LV., Gansa J., Morseb S.A., Pillaic S., Andersond K. and Hodged D.R., (2018). Evaluation of DNA Extraction Methods to Detect Bacterial Targets in Aerosol Samples. Journal of Microbiological Methods, Vol. 153, pp. 48–53. https://doi.org/10.1016/j.mimet.2018.09.006

24. Eisenhofer, R., Minich, J.J., Marotz, C., Cooper, A., Knight, R. and Weyrich, L.s., (2019). Contamination in Low Microbial Biomass Microbiome Studies: Issues and Recommendations. Trends in Microbiology, Vol. 27(2), pp. 105–117. https://doi.org/10.1016/j.tim.2018.11.003

25. Gill, C., van de Wijgert, J.H.H.M., Blow, F. and Darby, A.C., (2016). Evaluation of Lysis Methods for the Extraction of Bacterial DNA for Analysis of the Vaginal Microbiota. PLoS ONE, Vol. 11(9): e0163148. https://doi.org/10.1371/journal.pone.0163148

26. GraphPad Software, (2010). GraphPad Prism 5.04. San Diego California. https://www.graphpad.com/company Accessed January 2023

27. Grice E.A., Kong H.H., Conlan S., Deming C.B., Davis J., Young A.C., Bouffard G.G., Blakesley R.W., Murray P.R. and Green E.D., (2009). Topographical and temporal diversity of the human skin microbiome. Science, Vol. 324(5931), pp. 1190–1192. https://doi.org/10.1126/science.1171700

28. Gweon, H.S., Shaw, L.P., Swann, J., De Maio, N., AbuOun, M., Niehus, R., Hubbard, A.T.M., Bowes, M.J., Bailey, M.J., Peto, T.E.A., Hoosdally, S.J., Walker, A.S., P. Sebra, R.P., Crook, D.W., Anjum. M.F., Read, D.S., Stoesser, N., (2019). The impact of sequencing depth on the inferred taxonomic composition and AMR gene content of metagenomic samples. Environmental Microbiome, Vol.14(7). https://doi.org/10.1186/s40793-019-0347-1

29. Hammond, M., Homa, F., Andersson-Svahn, H., Ettema, T.J.G. and Joensson, H.N., (2016). Picodroplet partitioned whole genome amplification of low biomass samples preserves genomic diversity for metagenomic analysis. Microbiome, Vol. 4(52). https://doi.org/10.1186/s40168-016-0197-7

30. Jo, J.H., Kennedy, E.A. and Kong, H.H., (2016). Research techniques made simple: bacterial 16s ribosomal RNA gene sequencing in cutaneous research. Journal of Investigative Dermatology, Vol. 136(3), pp. e23–e27. https://doi.org/10.1016/j.jid.2016.01.005

31. Kang, DD, Froula, J., Egan, R. and Wang, Z., (2015). MetaBAT, an efficient tool for accurately reconstructing single genomes from complex microbial communities. PeerJ, 3:e1165. https://doi.org/10.7717/peerj.1165

32. Kong H.H., (2011). Skin microbiome: genomics-based insights into the diversity and role of skin microbes. Trends in Molecular Medicine, Vol. 17(6), pp. 320–328. https://doi.org/10.1016/j.molmed.2011.01.013

33. Kuczynski J., Lauber C.L., Walters W.A., Parfrey L.W., Clemente J.C., Gevers D. and Knight R., (2012). Experimental and analytical tools for studying the human microbiome. Nature Reviews Genetics, Vol. 13, pp.47–58. https://doi.org/10.1038/nrg3129

34. Li, D., Liu, C-M., Luo, R., Sadakane, K. and Lam, T-W., (2015). MEGAHIT: an ultra-fast single-node solution for large and complex metagenomics assembly via succinct de Bruijn graph. Bioinformatics, Vol. 31(10), pp. 1674–1676. https://doi.org/10.1093/bioinformatics/btv033

35. Liao, C-H and Shollenberger, L.M, (2003). Survivability and long-term preservation of bacteria in water and in phosphate-buffered saline. Letters in Applied Microbiology, Vol. 37(1), pp. 45-https://doi.org/10.1046/j.1472-765X.2003.01345.x

36. Liu Y-X., Qin Y., Chen T., Lu M., Qian X., Guo X. and Bai Y., (2020). Review: A practical guide to amplicon and metagenomic analysis of microbiome data. Protein Cell, Vol. 12(5), pp. 315–330. https://doi.org/10.1007/s13238-020-00724-8

37. Lou, Y.C., Hoff, J., Olm, M.R., West-Roberts, J., Diamond, S., Firek, B.A., Morowitz, M.J. and Banfield, J.F., (2022). Using strain-resolved analysis to identify contamination in metagenomics data. BioRxiv preprint. https://doi.org/10.1101/2022.01.16.476537

38. Lu J., Breitwieser, F.P., Thielen, P. and Salzberg, S.L., (2017). Bracken: estimating species abundance in metagenomics data. PeerJ Computer Science, 3:e104. https://doi.org/10.7717/peerj-cs.104

39. Maghini, D.G., Moss, E.L., Vance, S.E. and Bhatt, A.S., (2021). Improved high-molecular-weight DNA extraction, nanopore sequencing and metagenomic assembly from the human gut microbiome. Nature Protocols, Vol. 16(1), pp. 458–471. https://doi.org/10.1038/s41596-020-00424-x

40. Mandrekar P. V., Ma Z., Krueger S. and Cowan C., (2007). High-Concentration (>100ng/µl) Genomic DNA From Whole Blood Using the Maxwell® 16 Low Elution Volume Instrument. American Medical Association, Manual of Style, 10th edition. Promega Corporation Web site. https://www.promega.co.uk/resources/pubhub/high-concentration-genomic-dna-from-whole-blood-using-the-maxwell-16-low-elution-volume-instrumentUpdated2010. Accessed November 2020.

41. Marquet, M., Zöllkau, J., Pastuschek, J., Viehweger, A., Schleußner, E., Makarewicz, O., Pletz, M.W., Ehricht, R. and Brandt, C., (2022). Evaluation of microbiome enrichment and host DNA depletion in human vaginal samples using Oxford Nanopore’s adaptive sequencing. Scientific Reports, Vol. 12(4000). https://doi.org/10.1038/s41598-022-08003-8

42. Martzy, R., Bica-Schröder, K., Pálvölgyi, A.M., Kolm, C., Jakwerth, S., Kirschner, A.K.T., Sommer, R., Krska, R., Mach, R.L., Farnleitner, A.H. and Reischer, G.H., (2019). Simple lysis of bacterial cells for DNA-based diagnostics using hydrophilic ionic liquids. Scientific Reports, Vol. 9(13994). https://doi.org/10.1038/s41598-019-50246-5

43. Matthews TC, Bristow FR, Griffiths EJ, Petkau A, Adam J, Dooley D, Kruczkiewicz P, Curatcha J, Cabral J, Fornika D, Winsor GL, Courtot M, Bertelli C, Roudgar A, Feijao P, Mabon P, Enns E, Thiessen J, Keddy A, Isaac-Renton J, Gardy JL, Tang P, Consortium TI, Carrico JA, Chindelevitch L, Chauve C, Graham MR, McArthur AG, Taboada EN, Beiko RG, Brinkman FS, Hsiao WW, Domselaar GV. (2018). The Integrated Rapid Infectious Disease Analysis (IRIDA) Platform. bioRxiv preprint. https://doi.org/10.1101/381830

44. Moeller J.R., Moehn N.R., Waller D.M. and Givnish T.J., (2014). Paramagnetic Cellulose DNA Isolation Improves DNA Yield and Quality Among Diverse Plant Taxa. Applications in Plant Sciences, Vol. 2(10): pp. 1400048. https://doi.org/10.3732/apps.1400048

45. Narendran V., Visscher M.O., Abril I., Hendrix S.W. and Hoath S.B., (2010). Biomarkers of epidermal innate immunity in premature and fullterm infants. Pediatric Research, Vol. 67(4), pp. 382–386. https://doi.org/10.1203/PDR.0b013e3181d00b73

46. National Academies of Sciences, Engineering and Medicine (NASEM), (2018). Environmental Chemicals, the Human Microbiome, and Health Risk: A Research Strategy; National Academies Press: Washington, DC, USA. https://doi.org/10.17226/24960

47. Ogai K., Nagase S., Mukai K., Iuchi T., Mori Y., Matsue M., Sugitani K., Sugama J. and Okamoto S., (2018). A Comparison of Techniques for Collecting Skin Microbiome Samples: Swabbing Versus Tape-Stripping. Frontiers in Microbiology, Vol. 9(2362). https://doi.org/10.3389/fmicb.2018.02362

48. Olomu, I.N., Pena-Cortes, L.C., Long, R.A, Vyas, A., Krichevskiy, O., Luellwitz, R., Singh, P. and Mulks, M.H., (2020). Elimination of “kitome” and “splashome” contamination results in lack of detection of a unique placental microbiome. BMC Microbiology, Vol. 20(157). https://doi.org/10.1186/s12866-020-01839-y

49. Parks, D. H., Imelfort, M., Skennerton, C. T., Hugenholtz, P. and Tyson, G. W, (2015). CheckM: assessing the quality of microbial genomes recovered from isolates, single cells, and metagenomes. Genome Research, Vol.25(7), pp. 1043–1055. https://doi.org/10.1101/gr.186072.114

50. Pearman W.S., Freed N.E. and Silander O.K., (2020). Testing the advantages and disadvantages of short- and long-read eukaryotic metagenomics using simulated reads. BMC Bioinformatics, Vol. 21(220). https://doi.org/10.1186/s12859-020-3528-4

51. Phillips, S., Watt, R., Atkinson, T., Savva, G.M., Hayhoe, A. and Hall, L.J., (2021). The Pregnancy and EARly Life study (PEARL) - a longitudinal study to understand how gut microbes contribute to maintaining health during pregnancy and early life. BMC Pediatrics, Vol. 21(357). https://doi.org/10.1186/s12887-021-02835-5

52. Promega Maxwell® RSC Blood DNA Kit. https://www.promega.com/-/media/files/resources/protocols/technical-manuals/101/maxwell-rsc-blood-dna-kit-protocol.pdf?la=en Accessed November 2020.

53. RStudio Team (2021). RStudio: Integrated Development Environment for R. RStudio, PBC, Boston, MA. http://www.rstudio.com/ Accessed January 2023

54. Schindler, C.A. and Schuhardt, V.T., (1964). Lysostaphin: A New Bacteriolytic Agent for the Staphylococcus. Proceedings of the National Academy of Sciences of the United States of America, Vol. 51(3), pp. 414–421. https://doi.org/10.1073/pnas.51.3.414

55. Sczyrba, A., Hofmann, P., Belmann, P., et al., (2017). Critical assessment of metagenome interpretation—a benchmark of metagenomics software. Nature Methods, Vol. 14(11), pp. 1063–1071. https://doi.org/10.1038/nmeth.4458

56. Sfriso R., Egert M., Gempeler M., Voegeli R. and Campiche R., (2020). Revealing the secret life of skin - with the microbiome you never walk alone. International Journal of Cosmetic Science, Vol. 42(2), pp. 116–126. https://doi.org/10.1111/ics.12594

57. Singleton, C.M., Petriglieri1, F., Kristensen, J.M., Kirkegaard, R.H., Michaelsen, T.Y., Andersen, M.H., Kondrotaite, Z., Karst, S.M., Dueholm, M.S., Nielsen, P.H. and Albertsen, M., (2021). Connecting structure to function with the recovery of over 1000 high-quality metagenome-assembled genomes from activated sludge using long-read sequencing. Nature Communications, Vol. 12(2009). https://doi.org/10.1038/s41467-021-22203-2

58. Sui H., Weil A.A., Nuwagira E., Qadri F., Ryan E.T., Mezzari M.P., Phipatanakul W. and Lai P.S., (2020). Impact of DNA Extraction Method on Variation in Human and Built Environment Microbial Community and Functional Profiles Assessed by Shotgun Metagenomics Sequencing. Frontiers in Microbiology., Vol. 11(953). https://doi.org/10.3389/fmicb.2020.00953

59. The Human Microbiome Project Consortium, (2012). Structure, function and diversity of the healthy human microbiome. Nature, Vol. 486, pp. 207–214. https://doi.org/10.1038/nature11234

60. The Huttenhower Lab. Kneaddata. https://huttenhower.sph.harvard.edu/kneaddata/ Accessed January 2023

61. Toscano, M., De Grandi, R., Grossi, E. and Drago, L., (2017). Role of the Breast Milk-Associated Microbiota on the Newborns’ Immune System: A Mini Review. Frontiers in Microbiology, Vol. 8(2100). https://doi.org/10.3389/fmicb.2017.02100

62. Uritskiy, G.V., DiRuggiero, J. and Taylor, J., (2018). MetaWRAP - a flexible pipeline for genome-resolved metagenomic data analysis. Microbiome, Vol. 6(158). https://doi.org/10.1186/s40168-018-0541-1

63. Van Horn, K.G., Audette, C.D., Tucker, K.A. and Sebeck, D., (2008). Comparison of 3 swab transport systems for direct release and recovery of aerobic and anaerobic bacteria. Diagnostic Microbiology and Infectious Disease, Vol. 62(4), pp. 471–473. https://doi.org/10.1016/j.diagmicrobio.2008.08.004

64. Wang, Y., Zhao, Y., Bollas, A., Wang, Y. and Au, K.F., (2021). Nanopore sequencing technology, bioinformatics and applications. Nature Biotechnology, Vol. 39, pp. 1348–1365. https://doi.org/10.1038/s41587-021-01108-x

65. Wickham, H. 2009. ggplot2: Elegant Graphics for Data Analysis, Springer-Verlag New York, 1st Edition. https://doi.org/10.1007/978-0-387-98141-3

66. Wood, D.E., Lu, J. and Langmead, B., (2019). Improved metagenomic analysis with Kraken 2. Genome Biology, Vol.20(257). https://doi.org/10.1186/s13059-019-1891-0

67. Wu, Y-W., Simmons, B.A., Singer, S.W., (2016). MaxBin 2.0: an automated binning algorithm to recover genomes from multiple metagenomic datasets. Bioinformatics, Vol. 32(4), pp. 605–607. https://doi.org/10.1093/bioinformatics/btv638

68. Yan, W., Luo, B., Zhang, X., Ni, Y. and Tian, F., (2021). Association and Occurrence of Bifidobacterial Phylotypes Between Breast Milk and Fecal Microbiomes in Mother–Infant Dyads During the First 2 Years of Life. Frontiers in Microbiology, Vol. 12(669442). https://doi.org/10.3389/fmicb.2021.669442

69. Yuan, S., Cohen, D.B., Ravel, J., Abdo, Z., Forney, L.J., (2012). Evaluation of Methods for the Extraction and Purification of DNA from the Human Microbiome. PLoS ONE, Vol. 7(3), pp. e33865. https://doi.org/10.1371/journal.pone.0033865

70. Zhou, Q., Su, X. and Ning, K., (2014). Assessment of quality control approaches for metagenomic data analysis. Scientific Reports, Vol. 4(6957). https://doi.org/10.1038/srep06957

71. Zhu, T., Liu, X., Kong, F-Q., Duan, Y-Y., Yee, A.L, Kim, M., Galzote, C., Gilbert, J.A. and Quan, Z-X., (2019). Age and Mothers: Potent Influences of Children’s Skin Microbiota. Journal of Investigative Dermatology, 139(12), pp. 2497–505.e6. https://doi.org/10.1016/j.jid.2019.05.018

