## Supplementary Material for "An efficient method for high molecular weight bacterial DNA extraction suitable for shotgun metagenomics from skin swabs"

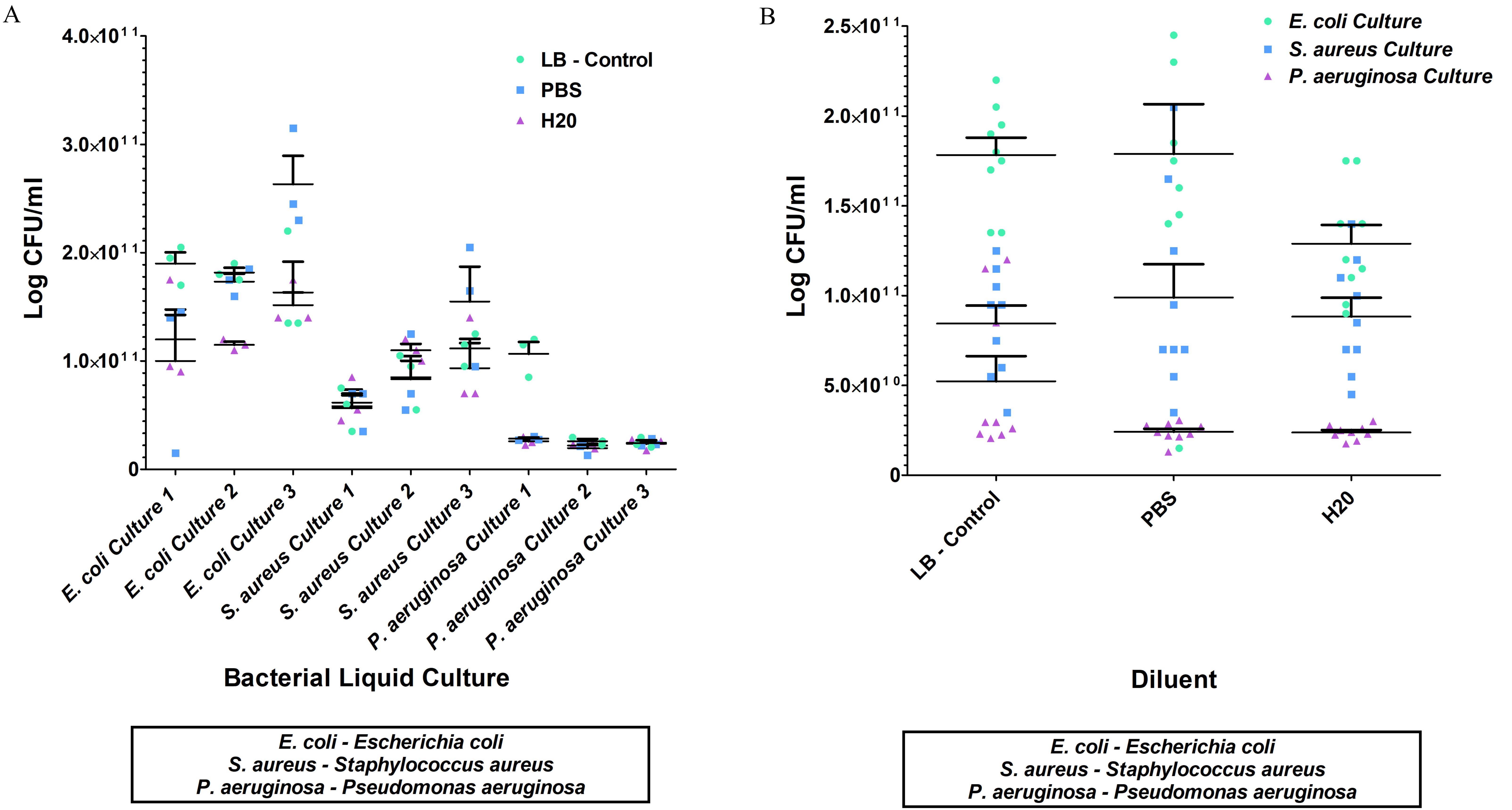


Figure S1 - Bacterial growth from cultures for *Escherichia coli*, *Staphylococcus aureus* and *Pseudomonas aeruginosa* that were processed and grown in 1x PBS and Milli-Q water. A and B: CFU/ml recovered for each isolate replicate cultured in LB broth, 1x PBS and Milli-Q water, and the reproducibility of replicates in each diluent condition. The horizontal bars on each plot show the average and vertical lines show the SEM.

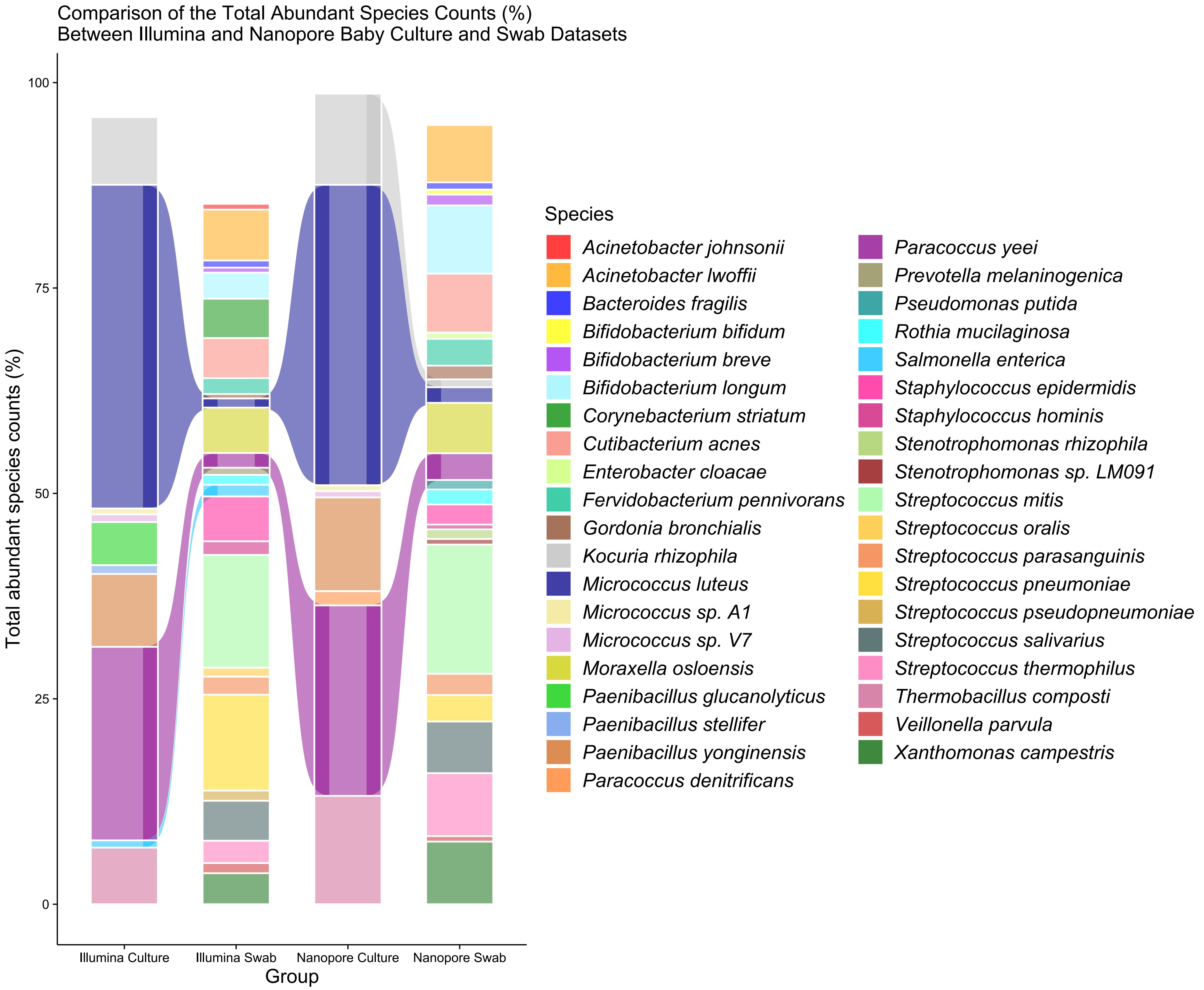


Figure S2 - Alluvial plot of the percentage total abundant species counts between Illumina and Nanopore SMS baby skin swab and culture datasets (one swab collected off one forearm from ten babies aged four months old). This was calculated by adding the total counts of each species, which were converted into a percentage of the total species count; only species with >0.5% of counts were used. The adjoining lines show the abundant skin species that were detected by both platforms.

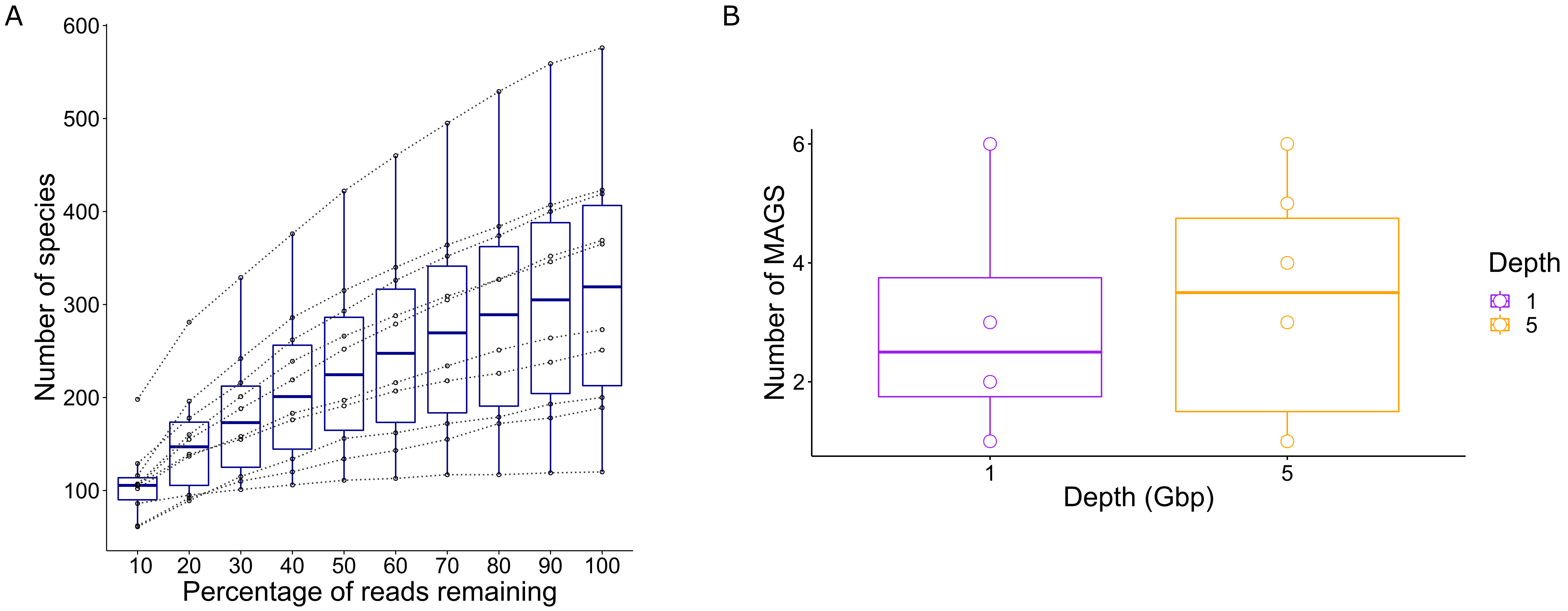


Figure S3 - Species and MAGs recovery from ten PEARL baby skin swabs (collected off one forearm from each baby aged four months old). A: Species recovery at 5Gbp sequencing depth and subsampled depths. The horizontal bars give the average number of species recovered at each level of reads, and the dots and lines show how each sample changes with reduced sequencing depth. 10%-100% reads represents 0.5Gbp-5Gbp at increments of 0.5Gbp. B: MAGs recovery at 1Gbp and 5Gbp sequencing depths. The horizontal bars give the average number of MAGs at each sequencing depth; the key to the right of the plot has colour coded the depth, with 1Gbp in purple and 5Gbp in orange. The vertical lines on each box plot show the SEM.

Table S1 – PEARL study baby metadata

| Sample ID | Age | Sex | Health status at birth | Breast fed at birth | Feed at week 3 | Feed at month 4 |
|---|---|---|---|---|---|---|
| Baby 1 | Month 4 | Female | Healthy | No | Missing Data | Missing Data |
| Baby 2 | Month 4 | Female | Healthy | No | Missing Data | Formula |
| Baby 3 | Month 4 | Male | Healthy | Yes | Breast | Breast |
| Baby 5 twin 1 | Month 4 | Female | Healthy | No | Breast/formula | Breast/formula |
| Baby 5 twin 2 | Month 4 | Male | Healthy | No | Breast/formula | Breast/formula |
| Baby 6 | Month 4 | Male | Healthy | Yes | Breast | Breast |
| Baby 7 | Month 4 | Female | Healthy | Yes | Breast | Breast |
| Baby 8 | Month 4 | Male | Healthy | Yes | Breast/formula | Formula |
| Baby 9 | Month 4 | Male | Healthy | No | Missing Data | Missing Data |
| Baby 10 | Month 4 | Male | Healthy | Breast/formula | Breast/formula | Formula |

Table S2 – Bacterial DNA concentrations from the six DNA extraction methods

| Overnight Liquid Culture | Method 1 BR Qubit (ng/µl) | Method 2 BR Qubit (ng/µl) | Method 3 BR Qubit (ng/µl) | Method 4 BR Qubit (ng/µl) | Method 5 BR Qubit (ng/µl) | Method 6 BR Qubit (ng/µl) |
|---|---|---|---|---|---|---|
| *E. coli* - Culture 1 | 19.9 | 11.5 | 36.8 | 47.2 | 40.9 | 46.8 |
| *E. coli* - Culture 2 | 33.2 | 11.3 | 46.5 | 55.3 | 54.9 | 38.5 |
| *P. aeruginosa* - Culture 1 | 8.68 | 18.6 | 63.9 | 30.8 | 88.4 | 96.7 |
| *P. aeruginosa* - Culture 2 | 7.96 | 12.2 | 33.5 | 42.7 | 97.7 | 104 |
| *S. aureus – Culture 1* | 9.7 | 5.18 | 32.8 | 29.7 | 51.2 | 37.8 |
| S. aureus – Culture 2 | 8.76 | 4.16 | 28.7 | 39.7 | 49.8 | 45.6 |

Table S3 – DNA concentrations of skin swabs

| Sample ID | HS Qubit (ng/µl) |
|---|---|
| Volunteer 1 – Swab 1 | < 0.50 |
| Volunteer 1 – Swab 2 | < 0.50 |
| Volunteer 2 – Swab 1 | < 0.50 |
| Volunteer 2 – Swab 2 | < 0.50 |
| Volunteer 3 – Swab 1 | < 0.50 |
| Volunteer 3 – Swab 2 | < 0.50 |
| Volunteer 4 – Swab 1 | < 0.50 |
| Volunteer 4 – Swab 2 | < 0.50 |
| Volunteer 5 – Swab 1 | 1.09 |
| Volunteer 5 – Swab 2 | 2.16 |
| Volunteer 6 – Swab 1 | 3.31 |
| Volunteer 6 – Swab 2 | 1.67 |
| Volunteer 7 – Swab 1 | 0.68 |
| Volunteer 7 – Swab 2 | 0.477 |
| Volunteer 8 – Swab 1 | < 0.50 |
| Volunteer 8 – Swab 2 | 0.156 |
| Volunteer 9 – Swab 1 | 0.837 |
| Volunteer 9 – Swab 2 | 0.809 |
| Volunteer 10 – Swab 1 | 6.52 |
| Volunteer 10 – Swab 2 | 9.73 |
| Volunteer 11 – Swab 1 | 10.5 |
| Volunteer 11 – Swab 2 | 5 |
| Volunteer 12 – Swab 1 | 1.69 |
| Volunteer 12 – Swab 2 | 3.63 |
| Positive Control 1 | 2.47 |
| Positive Control 2 | 1.1 |
| Positive Control 3 | 0.41 |
| Positive Control 4 | 0.261 |
| Negative Control | 0.064 |
| Baby 1 – Month 4 – Swab | 5.1 |
| Baby 2 – Month 4 – Swab | 4.05 |
| Baby 3 – Month 4 – Swab | 4.48 |
| Baby 5 – Twin 1 – Month 4 – Swab | 4.07 |
| Baby 5 – Twin 2 – Month 4 – Swab | 3.71 |
| Baby 6 – Month 4 – Swab | 4.47 |
| Baby 7 – Month 4 – Swab | 3.61 |
| Baby 8 – Month 4 – Swab | 3.91 |
| Baby 9 – Month 4 – Swab | 3.22 |
| Baby 10 – Month 4 – Swab | 3.46 |
| Positive Control (ATCC community) | 4.53 |
| Negative Control (PBS) | 2.69 |

Table S4 – MAGs recovered from ten skin swabs at 1Gbp and 5Gbp

| Sample | Sequence Data Generated (Gbp) | No. of MAGS | Genus |
|---|---|---|---|
| E003-B | 1 | 0 | N/A |
| E003-W1 | 1 | 0 | N/A |
| E005-1-B | 1 | 1 | *Paracoccus* |
| E005-2-B | 1 | 1 | *Bacillus* |
| E005-2-B | 1 | 1 | *Moraxella* |
| E005-2-B | 1 | 1 | *Pseudomonas* |
| E005-2-W3 | 1 | 0 | N/A |
| E005-2-M8 | 1 | 1 | *Streptococcus* |
| E005-2-M8 | 1 | 1 | *Neisseria* |
| E008-B | 1 | 0 | N/A |
| E014-B | 1 | 0 | N/A |
| E014-W3 | 1 | 0 | N/A |
| Positive Control | 1 | 1 | *Streptococcus* |
| Positive Control | 1 | 1 | *Micrococcus* |
| Positive Control | 1 | 1 | *Cutibacterium* |
| Positive Control | 1 | 1 | *Staphylococcus* |
| Positive Control | 1 | 1 | *Acinetobacter* |
| Positive Control | 1 | 1 | *Corynebacterium* |
| E003-B | 5 | 0 | N/A |
| E003-W1 | 5 | 0 | N/A |
| E005-1-B | 5 | 1 | *Paracoccus* |
| E005-1-B | 5 | 1 | *Paracoccus* |
| E005-1-B | 5 | 1 | *Paracoccus* |
| E005-2-B | 5 | 1 | *Moraxella* |
| E005-2-B | 5 | 1 | *Pseudomonas* |
| E005-2-B | 5 | 1 | *Bacillus* |
| E005-2-B | 5 | 1 | *Psychrobacter* |
| E005-2-W3 | 5 | 1 | *Cutibacterium* |
| E005-2-M8 | 5 | 1 | *Rothia* |
| E005-2-M8 | 5 | 1 | *Veillonella* |
| E005-2-M8 | 5 | 1 | *Neisseria* |
| E005-2-M8 | 5 | 1 | *Granulicatella* |
| E005-2-M8 | 5 | 1 | *Prevotella* |
| E008-B | 5 | 0 | N/A |
| E014-B | 5 | 1 | *Streptococcus* |
| E014-W3 | 5 | 0 | N/A |
| Positive Control | 5 | 1 | *Streptococcus* |
| Positive Control | 5 | 1 | *Staphylococcus* |
| Positive Control | 5 | 1 | *Corynebacterium* |
| Positive Control | 5 | 1 | *Cutibacterium* |
| Positive Control | 5 | 1 | *Acinetobacter* |
| Positive Control | 5 | 1 | *Micrococcus* |
